# Molecular basis for kinin selectivity and activation of the human bradykinin receptors

**DOI:** 10.1101/2021.05.27.446069

**Authors:** Yu-Ling Yin, Chenyu Ye, Fulai Zhou, Jia Wang, Dehua Yang, Wanchao Yin, Ming-Wei Wang, H. Eric Xu, Yi Jiang

## Abstract

Bradykinin and kallidin are endogenous kinin peptide hormones that belong to the kallikrein-kinin system and are essential to the regulation of blood pressure, inflammation, coagulation, and pain control. Des-Arg^10^-kallidin, the carboxy-terminal des-Arg metabolite of kallidin, and bradykinin selectively activate two G protein-coupled receptors, type 1 and type 2 bradykinin receptors (B1R and B2R), respectively. The hyperactivation of bradykinin receptors, termed “bradykinin storm”, is associated with pulmonary edema in COVID-19 patients, suggesting that bradykinin receptors are important targets for COVID-19 intervention. Here we report two G protein complex structures of B1R and B2R bound to des-Arg^10^-kallidin and bradykinin. Combined with functional analysis, our structures reveal the mechanism of ligand selectivity and specific activation of the bradykinin receptor. These findings also provide a framework for guiding drug design targeting bradykinin receptors for the treatment of inflammation, cardiovascular disorders, and COVID-19.

---

Kallikrein-kinin system (KKS) is a poorly understood hormonal system involved in the regulation of blood pressure, inflammation, coagulation, and pain control.^1–3^. The main components of KKS include the metabolic products of kinin peptides, such as bradykinin, kallidin (Lys-bradykinin) and its carboxy-terminal des-Arg metabolites, derived from different kininogen isoforms ^4^. These kinin peptides have highly conserved sequences with kallidin differing from bradykinin only by an additional N-terminal lysine, while des-Arg^10^-kallidin lacks the C-terminal Arg relative to kallidin. They are potent vasodilators and proinflammatory cytokines that activate bradykinin type 1 (B1R) and type 2 (B2R) receptors, two members of class A G protein-coupled receptors (GPCRs) ^5^.

Hyperactivation of KKS, termed “bradykinin storm”, was reported to be closely related to COVID-19 pathogenesis ^6^. Gene expression analyses of the bronchoalveolar lavage fluid from COVID-19 patients revealed a dramatic upregulation of B1R by ~3,000-fold and B2R by ~200 fold ^6^, respectively. The resulting “bradykinin storm” is thought to be responsible for most of the COVID-19 symptoms, including vascular leakage and pulmonary edema that are linked with hyperactivation of B1R and B2R ^7^. As such, blockade of B1R and B2R activation has been proposed as a therapeutic option to prevent acute respiratory distress syndrome in patients with COVID-19 ^8^.

B1R and B2R bind to kinin-derived peptide hormones and mediate transmembrane (TM) signaling primarily through G_q_ pathways. B2R is expressed in many normal tissues, whereas B1R expression is only induced in tissues under pathological conditions, such as inflammation ^1,9^. B1R and B2R share 34% identity in their amino acid sequences that are predicted to form a canonical GPCR fold of 7-TM helices, with a conserved peptide-binding pocket ^10^. Nevertheless, kinin peptides show different selectivity for bradykinin receptor subtypes. Specifically, bradykinin is one of the highest affinity kinin-derived peptides for B2R but exhibits low affinity for B1R, with over 10,000-fold selectivity for B1R ^11–13^. In contrast, des-Arg^10^-kallidin displays over 100,000-fold selectivity for B1R over B2R (Fig. 1a) ^11–13^. Extensive efforts have been made in defining the pharmacophore of antagonists and the molecular basis of ligand selectivity for kinins and other non-peptides using biochemical methods and molecular modeling ^14–19^. However, the underlined mechanisms for these peptide hormone-receptor subtype selectivity remain largely unknown due to the lack of structural evidence. Given their important physiological and pathological properties, it is of great value to elucidate molecular mechanisms for peptide recognition and bradykinin receptor activation. Here we report two cryo-electron microscopy (cryo-EM) structures of the B1R-G_q_ complex bound to des-Arg^10^-kallidin and the B2R-G_q_ complex bound to bradykinin. Combined with mutagenesis and functional analyses, our findings provide insights into specific recognition of kinin-derived peptide hormones by B1R and B2R and the molecular basis for receptor activation and G_q_ protein coupling.

**Fig. 1 │.**
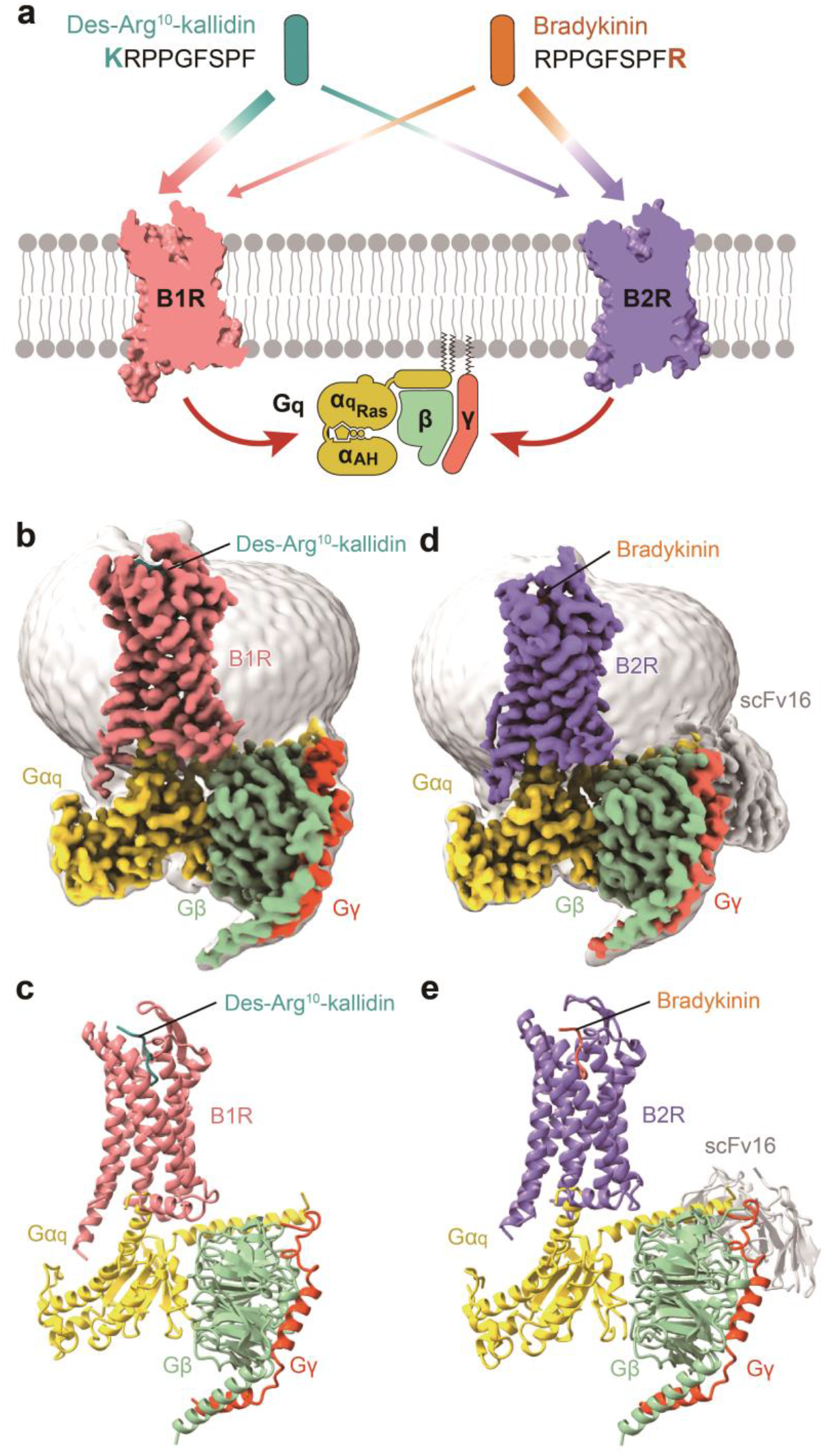
Cryo-EM structures of the des-Arg^10^-kallidin–B1R–G_q_ and bradykinin–B2R–G_q_ complexes. **a**, Schematic illustration of subtype selectivity for kinin and G_q_ protein-coupling of bradykinin receptors. Sequences of des-Arg^10^-kallidin and bradykinin are shown. **b, c**, Orthogonal views of the density map (**b**) and model (**c**) for the des-Arg^10^-kallidin–B1R–G_q_ complex. **d, e**, Orthogonal views of the density map (**d**) and model (**e**) for the bradykinin–B2R–G_q_ –scFv16 complex. Des-Arg^10^-kallidin is shown in cyan, des-Arg^10^-kallidin-bound B1R in salmon; bradykinin is displayed in orange, bradykinin-bound B2R in purple. The G_q_ heterotrimer is colored by subunits. Gα_q_, yellow; Gβ, pale green; Gγ, tomato; scFv16, grey.

## Results

### Structure determination of kinin-bound B1R and B2R

To stabilize B1R–G_q_ and B2R–G_q_ complexes, we applied the NanoBiT tethering method, a general strategy that has been used to obtain the structures of several GPCR-G protein complexes (Extended Data Fig. 1a, b) ^20–22^. An engineered Gα_q_ chimera was generated based on the mini-Gα_s/q_ 71 scaffold with its N-terminus replaced by corresponding sequences of Gα_i1_ to facilitate the binding of scFv16 (Extended Data Fig. 1c) ^23,24^. This analogous approach had been used in the structure determination of the 5-HT_2A_ R–G_q_ complex ^25^. Unless otherwise specified, G_q_ refers to G_q_ chimera used in the structure determination. Meanwhile, both B1R and B2R bear a tryptophan mutation at position 3.41 (F126^3.41^W for B1R and C146^3.41^W for B2R, superscripts refer to Ballesteros–Weinstein numbering ^26^), a known mutation that enhanced GPCR thermal stabilization ^27,28^. Both complexes were efficiently assembled on the membrane by co-expressing receptors with Gα_q_, Gβ1, and Gγ2 subunits (Extended Data Figs. 2a and 3a).

**Fig. 2 │.**
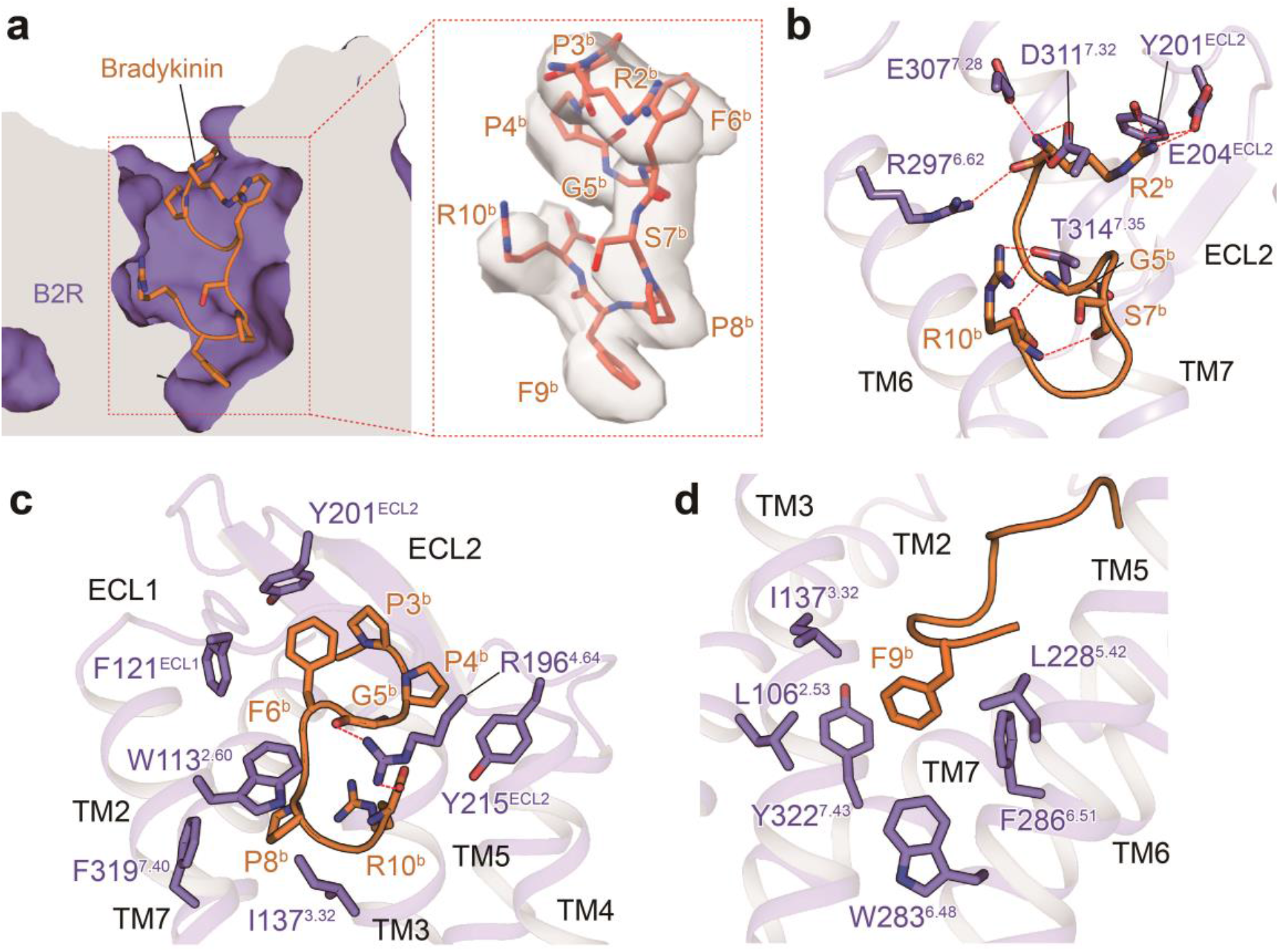
The bradykinin-binding pocket in B2R. **a**, Cross-section of the bradykinin-binding pocket in B2R. The cryo-EM density of bradykinin is highlighted. Side chains of the residues are displayed as sticks. Bradykinin is displayed in orange, B2R is colored in purple. **b-d**, Detailed interactions of bradykinin with residues in B2R. The binding site of R2^b^ and R10^b^ (**b**), P3^b^-P8^b^ (**c**), and F9^b^ (**d**) are shown. Hydrogen bonds and salt bridge are depicted as red dashed lines.

The structure of the des-Arg^10^-kallidin–B1R–G_q_ complex was defined with 633,636 final particles from 3,681,755 initial particles to a global nominal resolution of 3.0 Å (Fig. 1b, c and Extended Data Fig. 2 and Table 1). The structure of the bradykinin–B2R–G_q_ complex was determined with 664,416 final particles from 3,460,328 initial particles to a global nominal resolution of 2.9 Å (Fig. 1d, e and Extended Data Fig. 3 and Table 1). The overall conformation comparison shows highly similarity between two receptors, with an r.m.s.d. of 1.0 Å. Kinin peptides, TM bundles, extracellular loops (ECLs), and intracellular loops (ICLs) except ICL3 of both receptors show clear densities, enabling near-atomic modeling for two complexes. The majority of amino acid side chains were well-resolved in the refined final model (Fig. 1b-e and Extended Data Fig. 4). Thus, these two structures can provide detailed structural information of the peptide-binding pockets and receptor-G_q_ interaction interfaces.

**Fig. 3 │.**
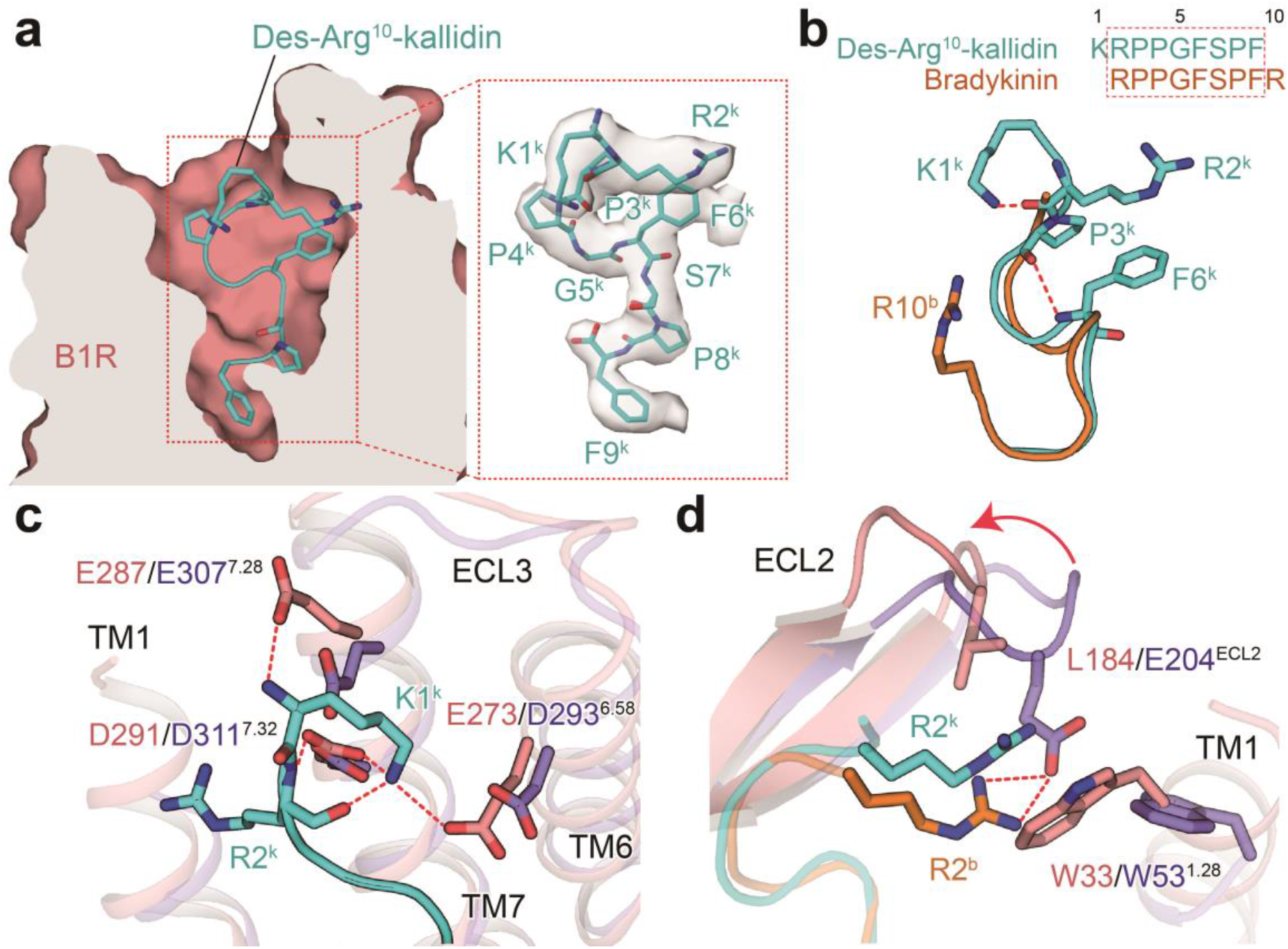
The des-Arg^10^-kallidin-binding pocket in B1R. **a**, Cross-section of the des-Arg^10^-kallidin-binding pocket in B1R. The cryo-EM density of des-Arg^10^-kallidin is highlighted. Side chains of the residues are displayed as sticks. **b**, A sequence and conformation comparison of des-Arg^10^-kallidin and bradykinin. Two intramolecular H-bonds are depicted as red dashed lines. Des-Arg^10^-kallidin is displayed in cyan, and bradykinin is colored in orange. **c**, Detailed interaction between K1^k^ and residues in B1R. **d**, Comparison of the binding mode of R2^b^ and R2^k^. The movement of ECL2 in B2R relative to that in B1R are highlighted in a red arrow. The salt bridges are shown as red dashed lines. Side chains of des-Arg^10^-kallidin and residues in two receptors are shown as sticks.

**Fig. 4 │.**
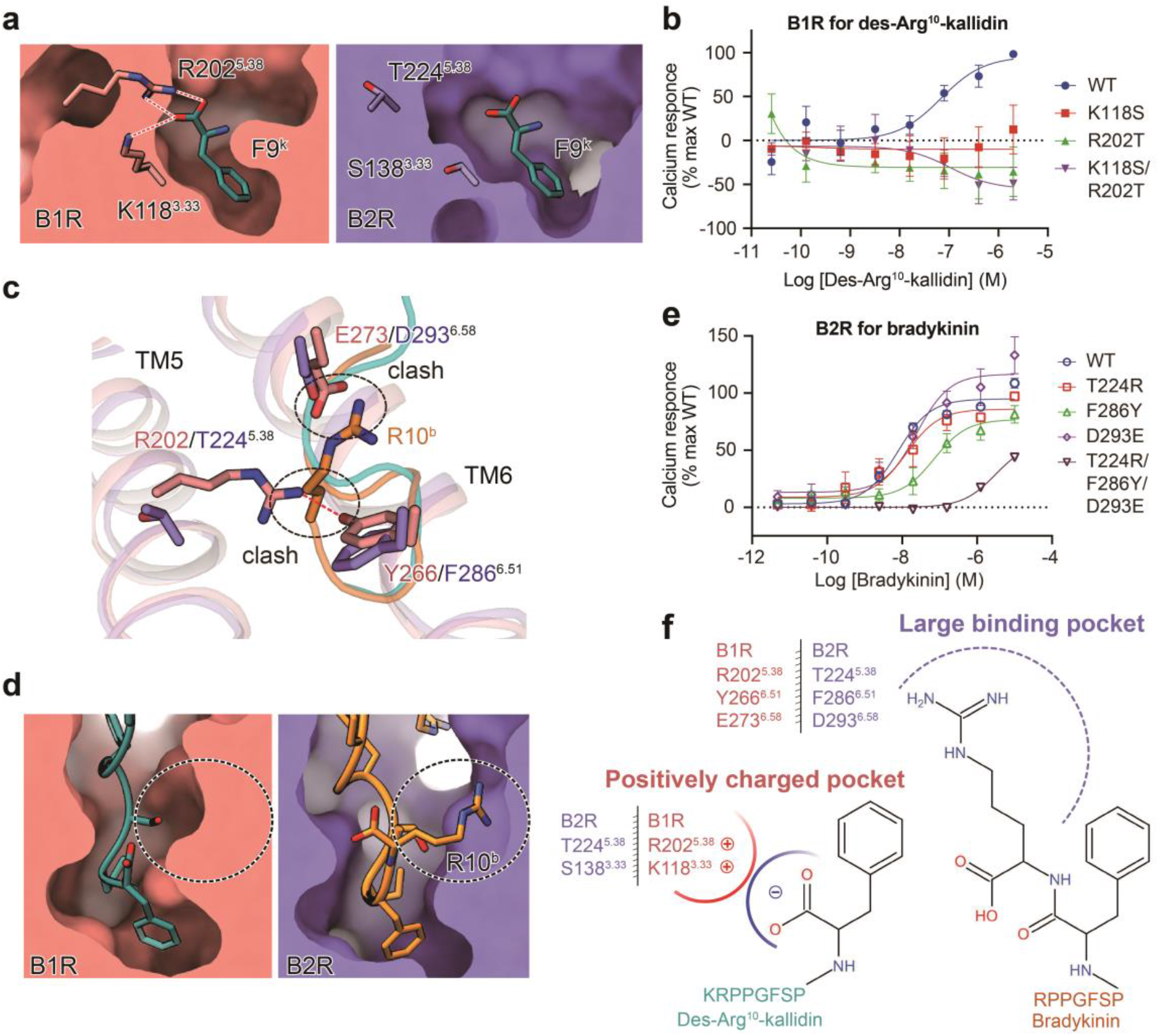
Molecular basis of kinin peptides selectivity for bradykinin receptors. **a**, Detailed interaction between F9^k^ and residues in B1R. F9^k^ and residues in B1R and the corresponding residues in B2R are shown as sticks. **b**, Effects of mutations in the F9^k^-binding pocket on calcium responses. Data are presented as means ± S.E.M. of three independent experiments. **c**, The binding site of R10^b^. Residues in the R10^b^-binding site in B2R and cognate residues in B1R are shown. The steric clash between side chain of R10^b^ and residues in B1R are highlighted as black dashed ovals. Polar interaction in **a** and **c** are shown as red dashed lines. **d**, A larger R10^b^ binding pocket in B2R relative to B1R. The pockets are highlighted as black dashed ovals. **e**, Effects of mutations in the R10^b^-binding pocket on calcium responses. **e**, Schematic model of the molecular basis of kinin peptides selectivity for bradykinin receptors. The chemical structures of F9^k^ in des-Arg^10^-kallidin and F9^b^ and R10^b^ in bradykinin are displayed. Residues in the positively charged pocket of B1R and the large binding pocket of B2R are highlighted. Data in **b** and **e** are displayed in Extended Data Table 3. Each data point presents mean ± S.E.M. of three independent experime nts. WT, wild-type.

### Molecular basis of bradykinin recognition by B2R

Bradykinin (RPPGFSPFR) occupies the orthosteric binding pocket comprising TM helices and ECLs except for TM1 and ECL3 (Extended Data Figs. 5 and 6). It presents an S-shaped overall conformation, with its C-terminus inserting deeply into the TMD core (Fig. 2a). This S-shaped fold is stabilized by two intramolecular H-bonds between the main chain of G5^b^ and R10^b^, as well as the backbone of S7^b^ and R10^b^ (Fig. 2b).

**Fig. 5 │.**
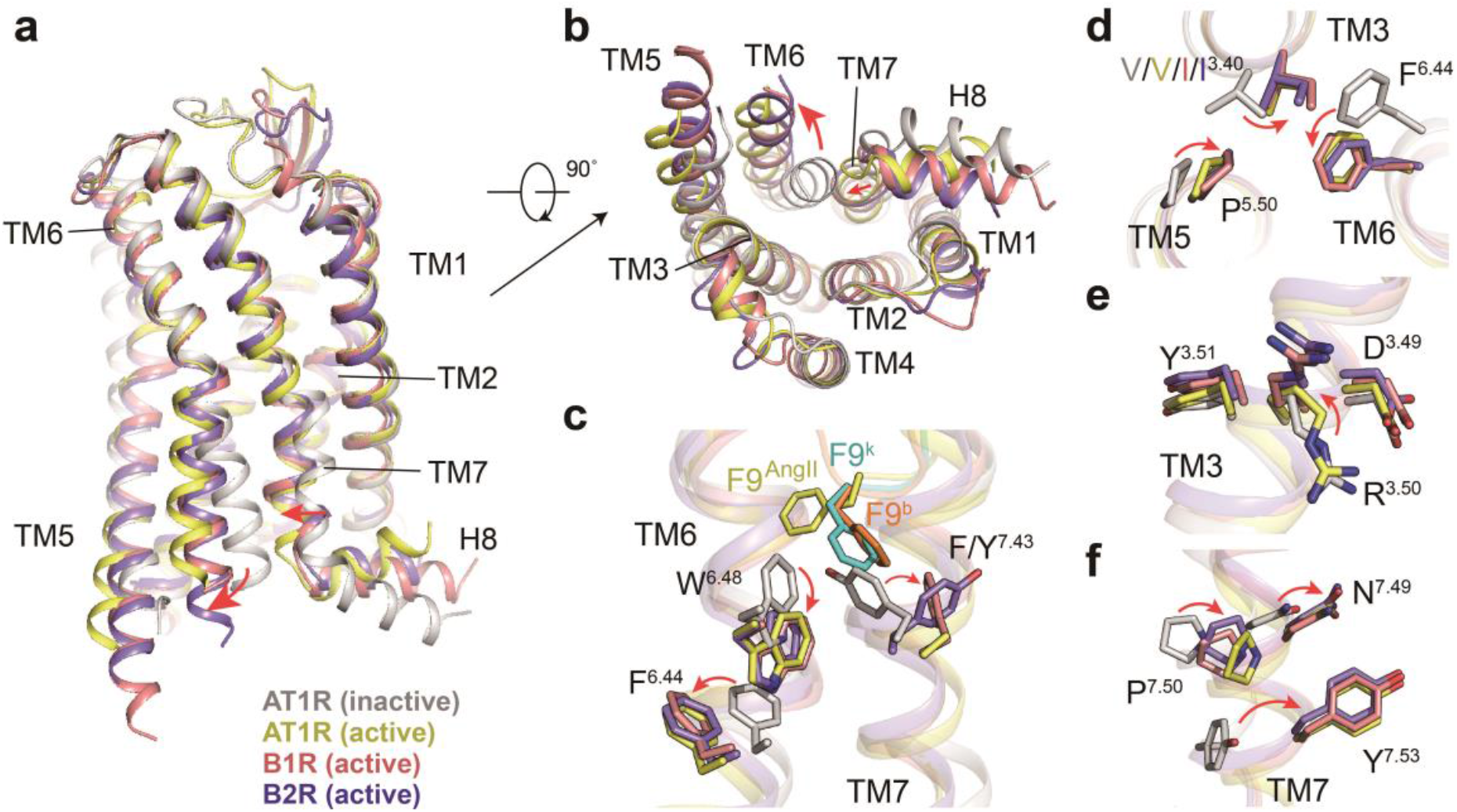
Activation mechanism of bradykinin receptors. **a, b**, Structural superposition of two active bradykinin receptors, inactive AT1R (PDB: 4YAY), and active AT1R (PDB: 6OS0) from the side (**a**) and cytoplasmic (**b**) views. The movement directions of TM6 and TM7 in bradykinin receptors relative to inactive AT1R are highlighted as red arrows. AT1R, angiotensin II receptor type 1. AT1R, B1R, and B2R are colored in grey, salmon, and purple, respectively. **c-f**, Conformational changes of the conserved “micro-switches” upon receptor activation, including Toggle switch (**c**), PIF (**d**), DRY (**e**), and NPxxY (**f**) motifs. F9^b^/F9^k^ triggered conformational changes of W^6.48^ and Y^7.43^ are highlighted. The conformational changes of residue side chains are shown as red arrows upon receptor activation. The complex structures were aligned by the receptors.

The sequence of bradykinin is featured by two positively charged arginines residing at both N- and C-terminus and the majority of hydrophobic amino acids at the middle segment of the peptide. The N-terminal R2 in bradykinin (refers to R2^b^) constitutes a stabilizing polar interaction network with ECL2, TM6, and TM7. The side chain of R2^b^ forms polar interactions with Y201^ECL2^ and E204^ECL2^. Its main chain NH group makes polar interactions with E307^7.28^ and D311^7.32^, while its backbone CO group builds an H-bond with R297^6.62^ (Fig. 2b). The C-terminal R10^b^ is also involved in polar interactions between bradykinin and B2R. Although the density of the guanidino group of R10^b^ is weak (Fig. 2a), it is indicative that the side chain of R10^b^ forms an H-bond with T314^7.35^, which is supported by diminished activity of bradykinin for B2R with T314^7.35^A mutation (Fig. 2b and Extended Data Fig. 7 and Table 2). Besides R2^b^ and R10^b^, the main chain CO group of G5^b^ makes an H-bond with R196^4.64^, which forms a salt bridge with the free carboxylic acid group of R10^b^ (Fig. 2c). The polar interaction network is essential for bradykinin-induced B2R activation since substituting R196^4.64^ with alanine entirely abolishes the activity of bradykinin (Extended Data Fig. 7 and Table 2).

P3^b^, P4^b^, F6^b^, P8^b^, and F9^b^ face hydrophobic environments within the B2R TMD pocket. P3^b^ and P4^b^ interact with the aromatic ring of Y201^ECL2^ and Y215^ECL2^, respectively (Fig. 2c). Y201^ECL2^, together with F121^ECL1^, make hydrophobic contact with F6^b^, which is also supported by the mutagenesis analysis (Fig. 2c and Extended Data Fig. 7 and Table 2). P8^b^ is surrounded by hydrophobic residues of TM2 (W113^2.60^), TM3 (I137^3.32^), and TM7 (F319^7.40^) (Fig. 2c). F9^b^ inserts deeply into a potent hydrophobic core comprised of residues in TM2 (L106^2.53^), TM3 (I137^3.32^), TM5 (L228^5.42^), TM6 (W283^6.48^ and F286^6.51^), and TM7 (Y322^7.43^) (Fig. 2d). Alanine mutations of these hydrophobic residues except for I137^3.32^ show a significant impact on bradykinin-induced B2R activation, indicating a potentially critical role of these hydrophobic residues near F9^b^ for bradykinin binding or B2R activation (Extended Data Fig. 7 and Table 2). Together, these detailed structural analyses provide important information to better understand the recognition mechanism of bradykinin by B2R.

### Molecular basis of des-Arg^10^-kallidin recognition by B1R

Des-Arg^10^-kallidin (KRPPGFSPF) shows high selectivity for B1R over B2R. Compared with bradykinin, des-Arg^10^-kallidin shares a conserved middle segment (RPPGFSPF) and sits in an almost identical orthosteric binding pocket with a similar S-shaped conformation (Fig. 3a and Extended Data Figs. 5 and 6). Nevertheless, distinct interactions are observed between two peptides and corresponding receptor subtypes, proving the basis for their receptor selectivity as described below.

In contrast to bradykinin, des-Arg^10^-kallidin has an additional lysine (K1^k^) at its N-terminus but lacks arginine that is located at the C-terminus of bradykinin (R10^b^) (Fig. 3b). Two extra intramolecular H-bonds exist between K1^k^ and backbone CO group of R2^k^, as well as backbone CO group of P3^k^ and NH group of F6^k^, causing a minor conformational change of des-Arg^10^-kallidin (Fig. 3b). The additional N-terminal K1^k^ forms polar interaction with E273^6.58^, E287^7.28^, and D291^7.32^ (Fig. 3c). Intriguingly, these residues are conserved in B2R (D293^6.58^, E307^7.28^, and D311^7.32^, Extended Data Fig. 6), which may explain the comparable B2R activation potency of Lys-bradykinin relative to bradykinin ^12^. Compared with R2^b^ in bradykinin, the equivalent R2^k^ in des-Arg^10^-kallidin presents distinct interactions with TM1 and ECL2. R2^k^ forms a cation-pi interaction with W33^1.28^ of B1R, while R2^b^ pushes W53^1.28^ away from the binding pocket owing to the steric hindrance (Fig. 3d). Additionally, ECL2 of B1R displays a smaller shift towards the peptide-binding pocket relative to B2R, which may be attributed to the lacking a corresponding salt bridge observed between R2^b^ and E204^ECL2^ in B2R (Fig. 3d).

### Molecular basis of kinin peptides selectivity for bradykinin receptors

Comparison of the binding modes between the two kinin peptides provides a framework for understanding kinin peptide selectivity by B1R and B2R. The free carboxylic acid backbone of F9^k^ engages a positively charged binding pocket and forms electrostatic interactions with K118^3.33^ and R202^5.38^ in B1R, which are not conserved in B2R (Fig. 4a). The cognate residues S138^3.33^ and T224^5.38^ in B2R fail to create a similar electrostatic environment, raising a hypothesis that the electrostatic pocket comprising of K118^3.33^ and R202^5.38^ is the determinant for selective binding of des-Arg^10^-kallidin to B1R over B2R. This hypothesis is supported by our mutagenesis studies that single or combined substitutions of K118^3.33^ and R202^5.38^ in B1R with serine and threonine, equivalent residues in B2R, abolished the activity of des-Arg^10^-kallidin (Fig. 4b, f and Extended Data Table 3). Our results are consistent with previous reports showing that K118^3.33^ attracts the negative charge of the C-terminus of B1-selective peptides and serves as a key residue in the selectivity of C-terminal des-Arg kinin peptides for B1R ^18,29^.

Compared with T224^5.38^, F286^6.51^, and D293^6.58^ in B2R, the cognate residues in B1R (R202^5.38^, Y266^6.51^, and E273^6.58^) are bulkier, resulting in insufficient space for interaction with the side chain of R10^b^ (Fig. 4c, d). The role of these residues in B2R selectivity for bradykinin is identified by swapping functional analysis. The triplicate swapping of T224^5.38^/F286^6.51^/D293^6.58^ with the cognate residues with larger side chains in B1R significantly impaired bradykinin activity (Fig. 4e and Extended Data Table 3). This finding suggests that a larger pocket comprised of T224^5.38^, F286^6.51^, and D293^6.58^ is crucial to bradykinin selectivity for B2R over B1R (Fig. 4f). Together, these data reveal the determinants of bradykinin receptor selectivity between bradykinin and des-Arg^10^-kallidin.

However, when mutating these kinin selectivity-related residues to cognate ones, only T224R mutation in B2R showed slightly increased activity of des-Arg^10^-kallidin. Other residue substitutions did not cause significantly increased activities of bradykinin and des-Arg^10^-kallidin for B1R and B2R, respectively (Extended Data Fig. 7i, j). It seems that only swapping the residues in the electrostatic pocket in B1R or a larger pocket in B2R failed to make the two receptors possess high-affinity for kinins. Thus, we believe that the residues in these two pockets are not entirely responsible for kinin selectivity.

### Activation mechanism of B1R and B2R

A structural comparison of B1R and B2R complexes to their closely related angiotensin II receptor type 1 (AT1R) in the inactive (PDB: 4YAY ^30^) and active states (PDB: 6OS0 ^31^) sheds light on the basis of bradykinin receptor activation. The structural comparison demonstrates that both B1R and B2R adopt fully active conformations similar to the active AT1R (Fig. 5a). Compared with the inactive AT1R, they show a remarkable outward displacement of the cytoplasmic end of TM6, a hallmark of class A GPCR activation, along with an inward movement of TM7 cytoplasmic end (Fig. 5a, b) ^32^.

Although bradykinin and des-Arg^10^-kallidin present different binding selectivity, they may activate bradykinin receptors through a common mechanism. The side chains of F9^b^ and F9^k^ insert into a conserved hydrophobic crevice at the bottom of the peptide-binding pocket and trigger rotameric switch of W^6.48^, the toggle switch residue, which further facilitates the swing of F^6.44^ and initiates the rotation of TM6 (Fig. 5c). Meanwhile, the steric clash between F9^k^/F9^b^ and F/Y^7.43^ would drive the latter swinging away from the receptor helical core and the inward shifting of the cytoplasmic end of TM7 (Fig. 5c). [Leu^9^, des-Arg^10^]kallidin, in which F9^k^ of des-Arg^10^-kallidin is substituted with a smaller bulky amino acid (leucine), loses its agonistic activity with conversion to a B1R antagonist, supporting the critical role of F9^k^ in B1R activation ^33,34^. The switches of W^6.48^ and F/Y^7.43^ further trigger the active-like conformational changes of “micro-switch” residues (toggle switch W^6.48^ and PIF, DRY, and NPxxY motifs), leading to an agonism signal transduction to the cytoplasmic end of the receptor (Fig. 5c-f).

Structural comparison of B1R and B2R with their closely related class A GPCR member AT1R in the active state (PDB: 6OS0 ^31^) suggests a common mechanism of receptor activation. The bound endogenous peptide hormones des-Arg^10^-kallidin, bradykinin, and angiotensin II share conserved C-terminal phenylalanine, which inserts into the peptide-binding pockets of corresponding receptors at a comparable depth (Extended Data Fig. 8). Moreover, although differing in side chain orientations, these phenylalanines are buried within a similar hydrophobic environment, indicating a universal activation mechanism of these closely related GPCRs (Extended Data Fig. 8b).

Structural superposition of G_q_-coupled B1R and B2R complexes with the G_q_-coupled 5-HT_2A_ R (PDB: 6WHA ^25^) and G_11_-coupled M1R (PDB: 6OIJ ^35^) by receptors shows nearly identical conformations of TM6 and TM7 (Extended Data Fig. 9a), suggesting that G_q_-coupled receptors have the similar conformation to G_11_-coupled receptors. In addition, compared with these two G_q/11_-coupled receptors, the helix 8 of B1R and B2R is closer to the Gβ subunit, which may be attributed to the intramolecular salt bridge formed between K^8.53^ and E^2.40^ (Extended Data Fig. 9b). On the G protein side, the α5 helix of Gα_q_ shifts 4 Å for both B1R and B2R compared with that of G_11_-coupled M1R (measured at Cα of Y^H5.23^) and moves half-helical turn upward towards the cytoplasmic cavity of the receptor (2 Å for B1R and 3 Å for B2R, respectively) relative to G_q_-coupled 5-HT_2A_ receptor. Meanwhile, the Gα_q_ N-termini of B1R and B2R undergo notable shifts seen across the G_q/11_-coupled class A GPCRs (Extended Data Fig. 9c).

## Discussion

Bradykinin receptors are involved in various clinical symptoms and their use as therapeutic targets remains the focus of extensive investigations. Recently, decoding of the bradykinin inflammatory pathway in COVID-19, known as “bradykinin storm”, highlights the implication of bradykinin receptor modulators as a potential treatment of COVID-19. In this study, we determined two G_q_-coupled structures of the human B1R and B2R bound to selective kinin peptides, des-Arg^10^-kallidin and bradykinin, respectively. In combination with functional analyses, these structures enhance our understanding of the molecular basis of kinin peptides recognition and activation of B1R and B2R.

Intriguingly, it has been recently predicted that des-Arg^10^-kallidin and bradykinin show distinct V-and S-shaped conformations, respectively. These distinct conformations result in different presentations of kinin peptides’ N-and C-termini toward their receptors ^18^. In our structural model, bradykinin adopts an overall similar S-shaped conformation but significantly differs in the orientation of the C-terminal charged arginine (R10^b^), with an overall r.m.s.d. of 1.8 Å (Extended Data Fig. 10a). The side chain of R10^b^ sits in the gap between TM6 and TM7 and forms a salt bridge with T314^7.35^. In contrast, R10^b^ points to TM5 and forms a salt bridge with E221^5.35^ in the predicted model. It is worth noting that des-Arg^10^-kallidin in the structural model shows an entirely different conformation, presenting an S-shaped but not a V-shaped fold with an r.m.s.d. of 3.4 Å (Extended Data Fig. 10b). Even so, the middle segment of des-Arg^10^-kallidin (P3^k^-F6^k^) in both our structure and NMR model displays a similar β-turn-like conformation, which might be stabilized by the intramolecular H-bond made by backbone NH of F6^k^ and CO of P3^k^, as observed in B1R complex structure. A des-Arg^10^-kallidin analog with the methylated amide of F6^k^, which may disturb this H-bond and central β-turn, showed a 1,000-fold lower binding affinity for B1R than the native peptide, indicating the importance of this intramolecular interaction in maintaining conformation stability of des-Arg^10^-kallidin ^18^. It was predicted that the C-terminal segment of bradykinin (S7^b^-R10^b^) adopted a β-turn conformation, which might be one of the necessities for high affinity to B2R ^17^. The β-turn constitutes a molecular basis for designing B2R ligands, including the only approved B2R antagonist icatibant ^16,17^. Consistently, we also observed a similar β-turn conformation, which is stabilized by two intramolecular H-bonds in B2R structure. This β-turn forces F9^b^ inserting deeply into the TMD core and engaging with hydrophobic residues at the bottom of the B2R pocket. Our alanine mutagenesis analysis on these hydrophobic residues further supports a potential role of the β-turn conformation of bradykinin in B2R activation.

Bradykinin receptors exhibit exquisite selectivity for bradykinin and des-Arg^10^-kallidin, and our findings provide a framework to depict the subtype selectivity of kinin peptides for B1R and B2R. It was found that the residue environments surrounding C-termini amino acids of bradykinin and des-Arg^10^-kallidin are critical to such a selectivity. K118^3.33^ and R202^5.38^, which constitute a positively charged pocket interacting with the free carboxylic acid group of F9^k^, are determinants of the preference of des-Arg^10^-kallidin to B1R. Additionally, the H-bond between R202^5.38^ and Y266^6.51^ in B1R creates an inaccessible space for bradykinin. Meanwhile, the smaller side chains of T224^5.38^, F286^6.51^, and D293^6.58^ in B2R relative to equivalent residues in B1R, create a larger pocket space to accommodate R10^b^, thereby revealing the molecular basis of the higher B2R selectivity by bradykinin. Coincidentally, the NMR model predicted the same H-bond between R202^5.38^ and Y266^6.5118^. A molecular modeling study on B1R and B2R also speculated that the smaller size of T^5.38^ in B2R relative to cognate residue R^5.38^ in B1R allows non-peptide antagonists to access an aromatic pocket composed of W^6.48^, F^6.51^, and Y^7.43^, which is inaccessible for B1R ^15,16,19^. However, R/T^5.38^ in B1R and B2R cannot hamper the engagement of kinins with this aromatic pocket, which accommodates F9^k^/F9^b^ in our structures. Conversely, R/T^5.38^ is involved in the binding of R10^b^ to a larger pocket in B2R, which partially determines bradykinin selectivity for B2R over B1R. Additionally, we further propose a common activation mechanism for B1R and B2R, through structural comparison with AT1R. With an in-depth knowledge about ligand selectivity and receptor activation, new opportunities will arise to design potent and efficacious modulators of B1R and B2R for the treatment of inflammation, cardiovascular disorders, and COVID-19.

## Acknowledgements

The Cryo-EM data of bradykinin–B2R–G_q_ complex were collected at Cryo-Electron Microscopy Research Center, Shanghai Institute of Material Medica. Cryo-EM data collection of des-Arg^10^-kallidin–B1R–G_q_ was carried out at the Shuimu BioSciences Ltd. We thank all the staff at these two cryo-EM facilities for their technical support. This work was partially supported by the Ministry of Science and Technology (China) grants 2018YFA0507002 (H.E.X.) and 2018YFA0507000 (M.-W.W.); National Natural Science Foundation of China 31770796 (Y.J.), 81872915 (M.-W.W.), 82073904 (M.-W.W.), 81773792 (D.Y.), and 81973373 (D.Y.); National Science and Technology Major Project of China – Key New Drug Creation and Manufacturing Program 2018ZX09711002-002-002 (Y.J.), 2018ZX09735–001 (M.-W.W.), and 2018ZX09711002–002–005 (D.Y.); the Shanghai Municipal Science and Technology Commission Major Project 2019SHZDZX02 (H.E.X.); the CAS Strategic Priority Research Program XDB37030103 (H.E.X.); Novo Nordisk-CAS Research Fund grant NNCAS-2017–1-CC (D.Y.); and the Young Innovator Association of CAS (2021278 to W.Y.).

## Author Contributions

Y.-L.Y. screened the expression constructs, optimized the bradykinin-Gq protein complexes, prepared the protein samples for final structure determination, participated in cryo-EM grid inspection, data collection, and model building; C.Y. and J.W. designed the mutations and executed the functional studies; F.Z. build and refine the structure models; M.-W.W and D.Y. supervised functional assay development and data analysis; W.Y. designed G_q_ protein constructs and prepared samples for the cryo-EM; H.E.X. and Y.J. conceived and supervised the project and initiated collaborations with M.-W.W.; Y.J. and Y.-L.Y. prepared the figures and drafted manuscript; Y.J., H.E.X., and M.-W.W wrote the manuscript with inputs from all authors.

## Data availability

All data is available in the main text or the supplementary materials. Materials are available from the corresponding authors upon reasonable request.

---

**Extended Data Figure. 1│.**
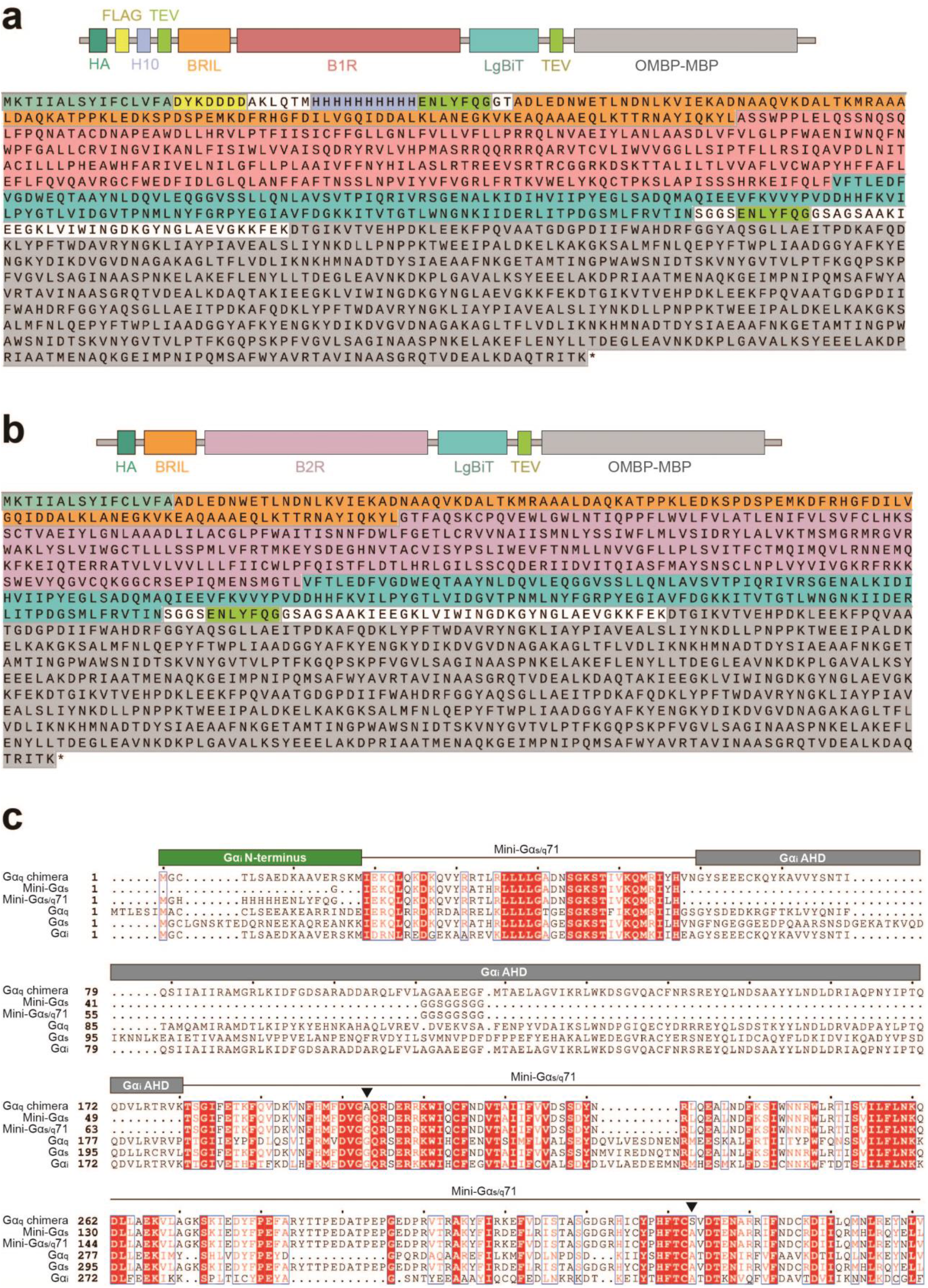
B1R, B2R and Gα_q_ chimera/mutant constructs used in this study. **a, b**, Schematic representation and amino acid sequences of B1R (**a**) and B2R (**b**) constructs. The sequences of different components in two constructs are shown in indicated colors. **c**, Sequence alignment of Gα_q_ chimera with wild-type and other engineered Gα subunits. The skeleton is based on mini-Gα_s/q_ 71 ^23^. N-terminus is replaced by Gα_i1_ for scFv16 binding (green). The α-helical domain (AHD) is substituted by the corresponding sequence of the human Gα_i_ subunit (grey). Two dominant-negative mutations are indicated by black triangles. Mini-Gα_s_ is the engineered Gα_s_ used for the structure determination of A_2A_ R–mini-G_s_ complex (PDB: 5G53 ^36^).

**Extended Data Figure. 2│.**
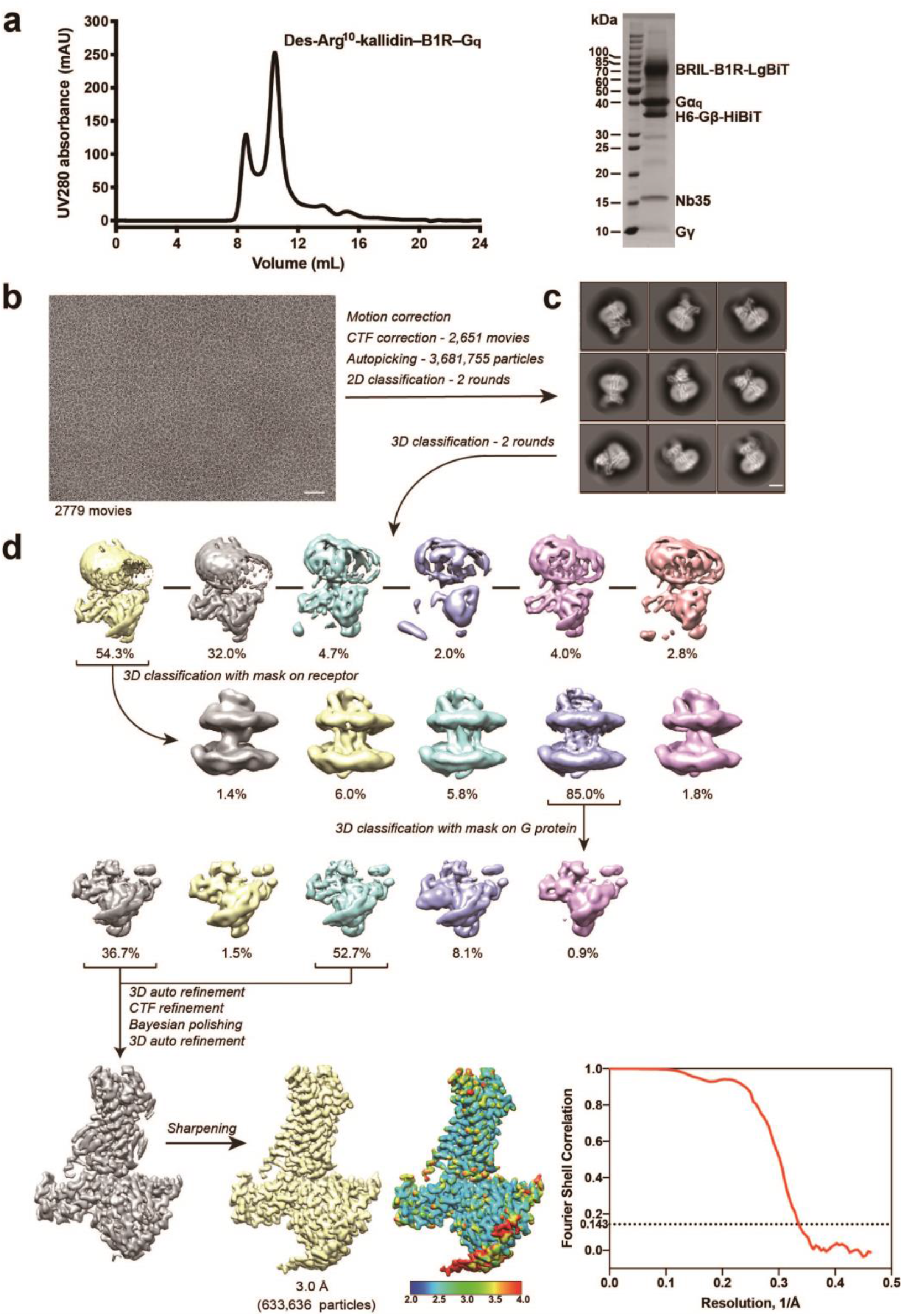
Des-Arg^10^-kallidin–B1R–G_q_ complex purification and cryo-EM data processing. **a**, Representative size-exclusion chromatography elution profile and SDS-PAGE analysis of the des-Arg^10^-kallidin–B1R–G_q_ complex. **b**, Cryo-EM micrograph of the des-Arg^10^-kallidin–B1R–G_q_ complex. Scale bar, 50 nm. **c**, Representative 2D average classes of the des-Arg^10^-kallidin–B1R–G_q_ complex. Scale bar, 5 nm. **d**, Flowchart of cryo-EM data processing.

**Extended Data Figure. 3│.**
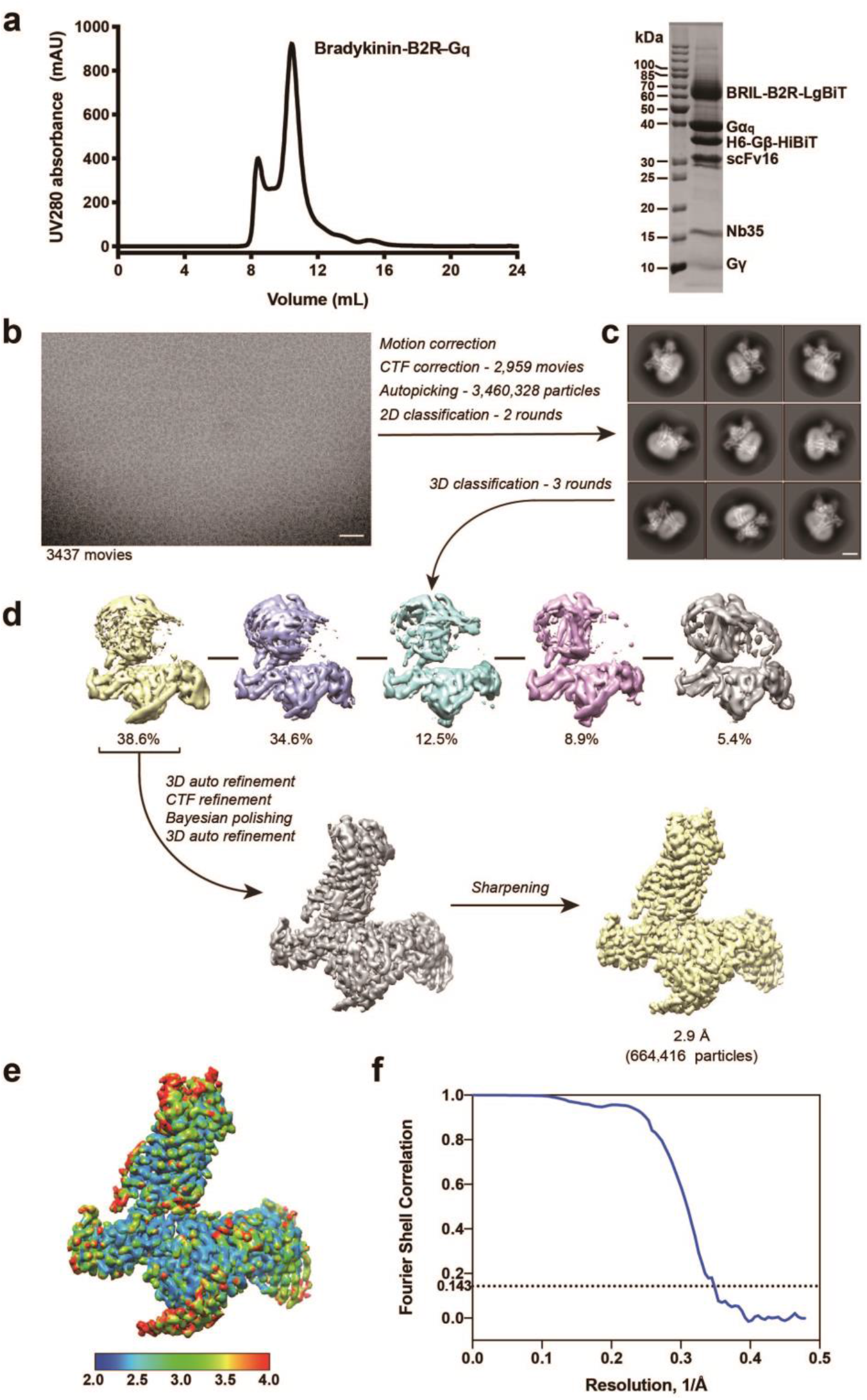
Bradykinin–B2R–G_q_ complex purification and cryo-EM data processing. **a**, Representative size-exclusion chromatography elution profile and SDS-PAGE analysis of the bradykinin–B2R–G_q_ complex. **b**, Cryo-EM micrograph of the bradykinin–B2R–G_q_ complex. Scale bar, 50 nm. **c**, Representative 2D average classes of the bradykinin–B2R–G_q_ complex. Scale bar, 5 nm. **d**, Flowchart of cryo-EM data processing. **e**, Cryo-EM map of the bradykinin–B2R–G_q_ complex, colored by local resolution (Å) calculated using Resmap package. **f**, “Gold-standard” FSC curves.

**Extended Data Figure. 4│.**
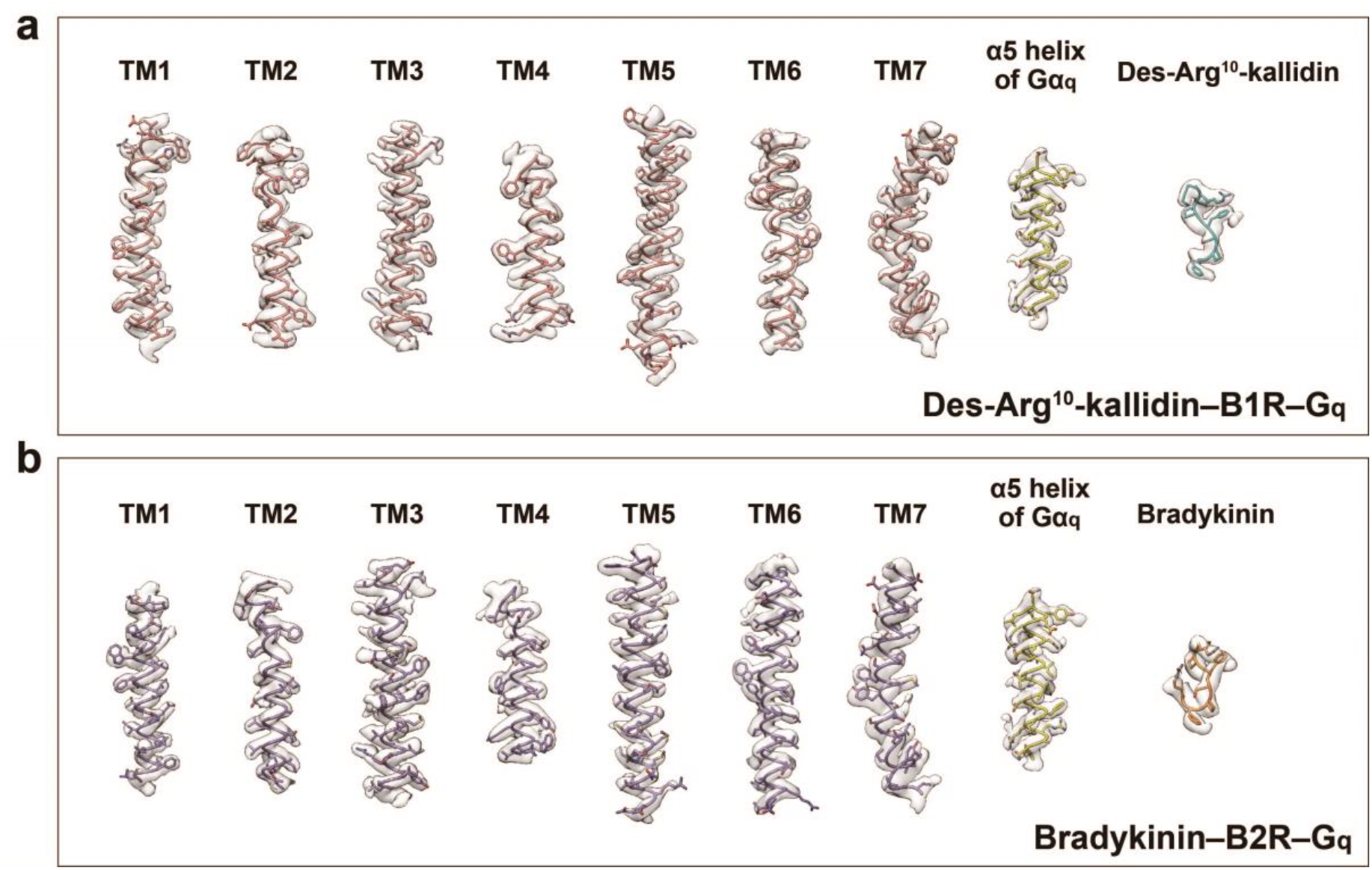
Cryo-EM density maps of the des-Arg^10^-kallidin–B1R–G_q_ and bradykinin–B2R–G_q_ complex. **a**, Cryo-EM density maps of the seven transmembrane (TM) helices of B1R, α5 helix of Gα_q_, and des-Arg^10^-kallidin. **b**, Cryo-EM density maps of the seven TM helices of B2R, α5 helix of Gα_q_, and bradykinin.

**Extended Data Figure. 5│.**
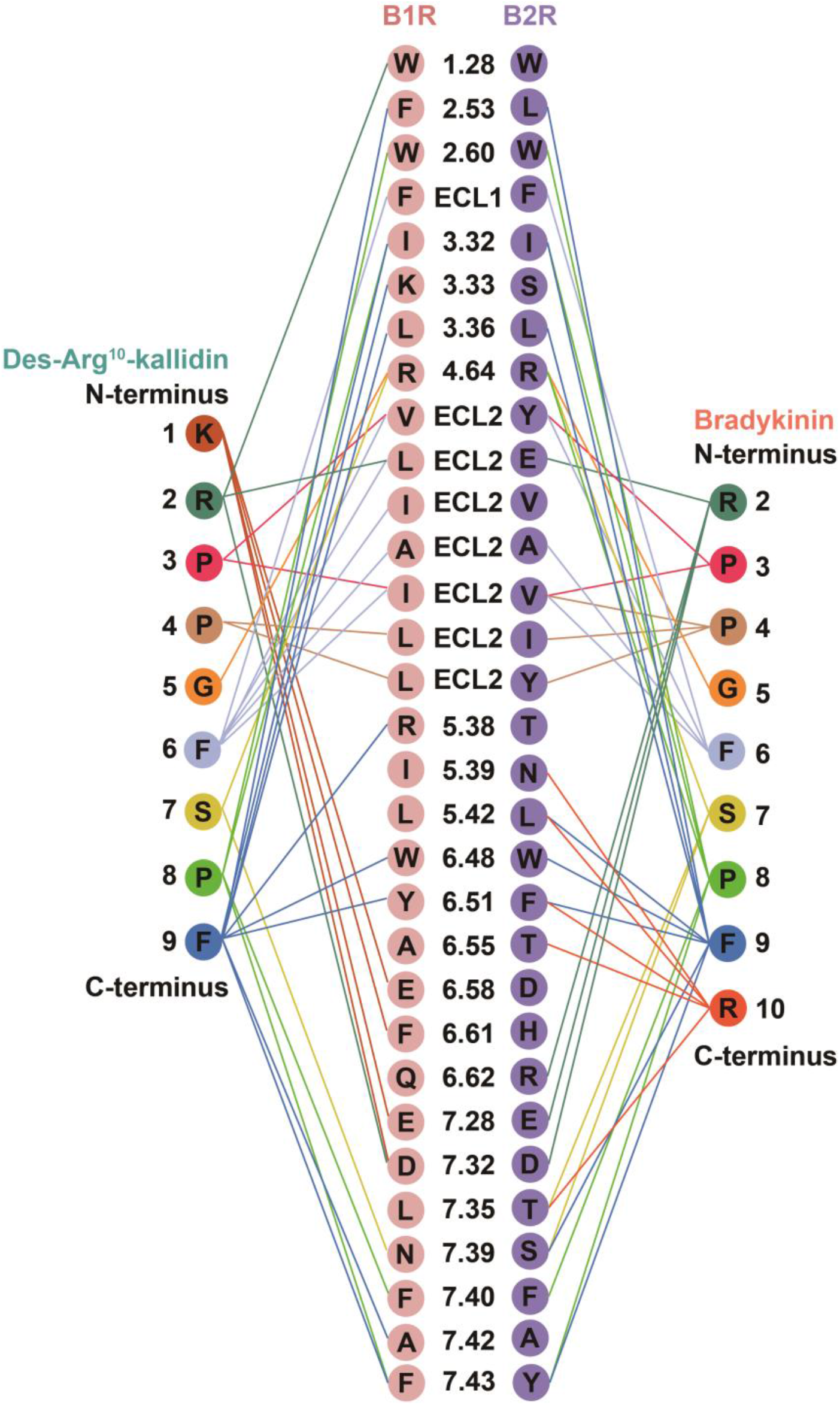
Representative interaction network of des-Arg^10^-kallidin bound to B1R and bradykinin bound to B2R.

**Extended Data Figure. 6│.**
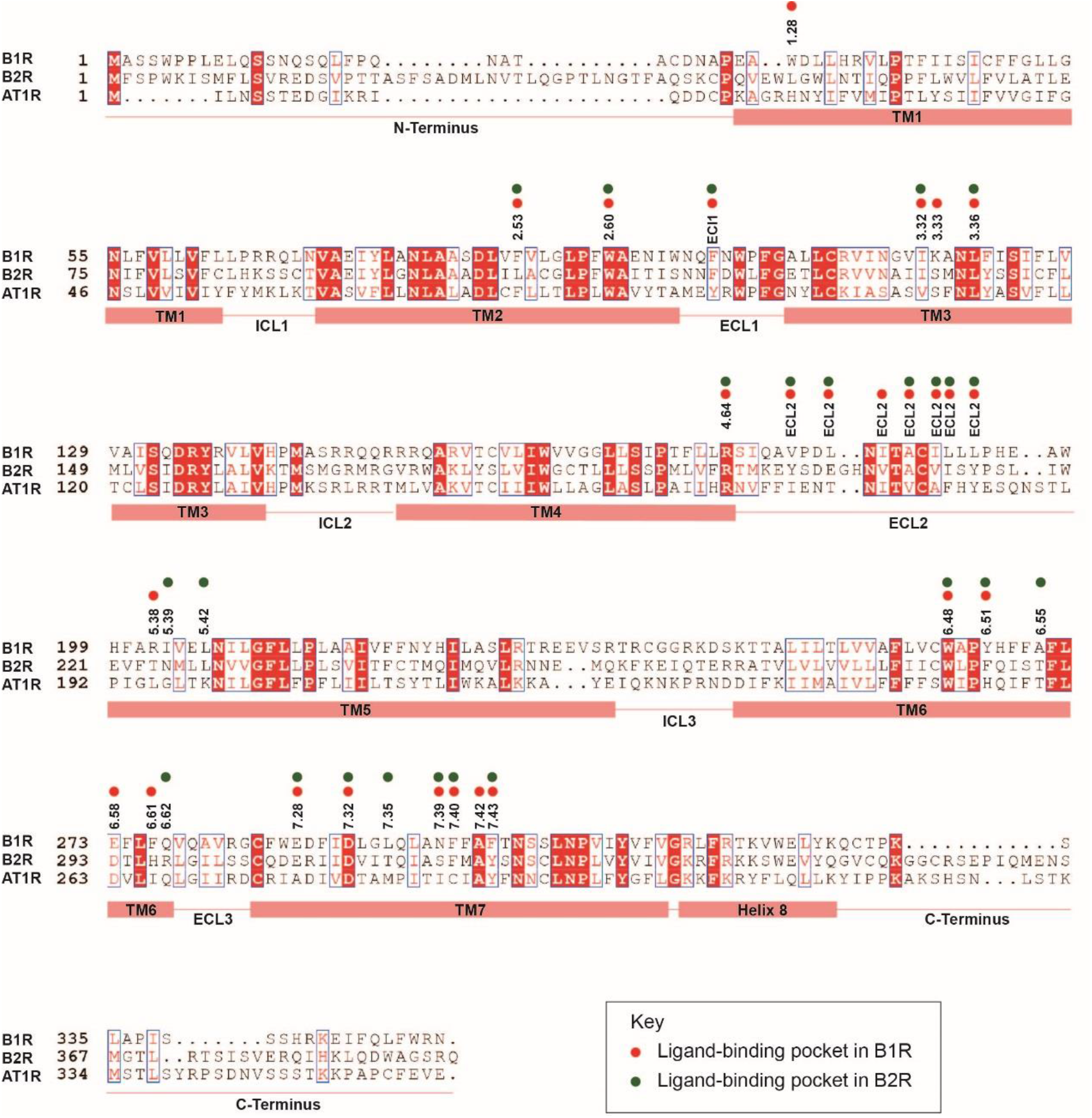
Sequence alignment of the bradykinin/angiotensin receptor subfamily. The sequence alignment of B1R, B2R, and angiotensin II receptor type 1 (AT1) was generated using NCBI and the graphics was created on the ESPript 3.0 server. α-helices for B1R are shown as columns underneath the sequences. Red dots represent the binding-pocket residues of B1R bound to des-Arg^10^-kallidin, and green dots represent the binding-pocket residues of B2R bound to bradykinin.

**Extended Data Figure. 7│.**
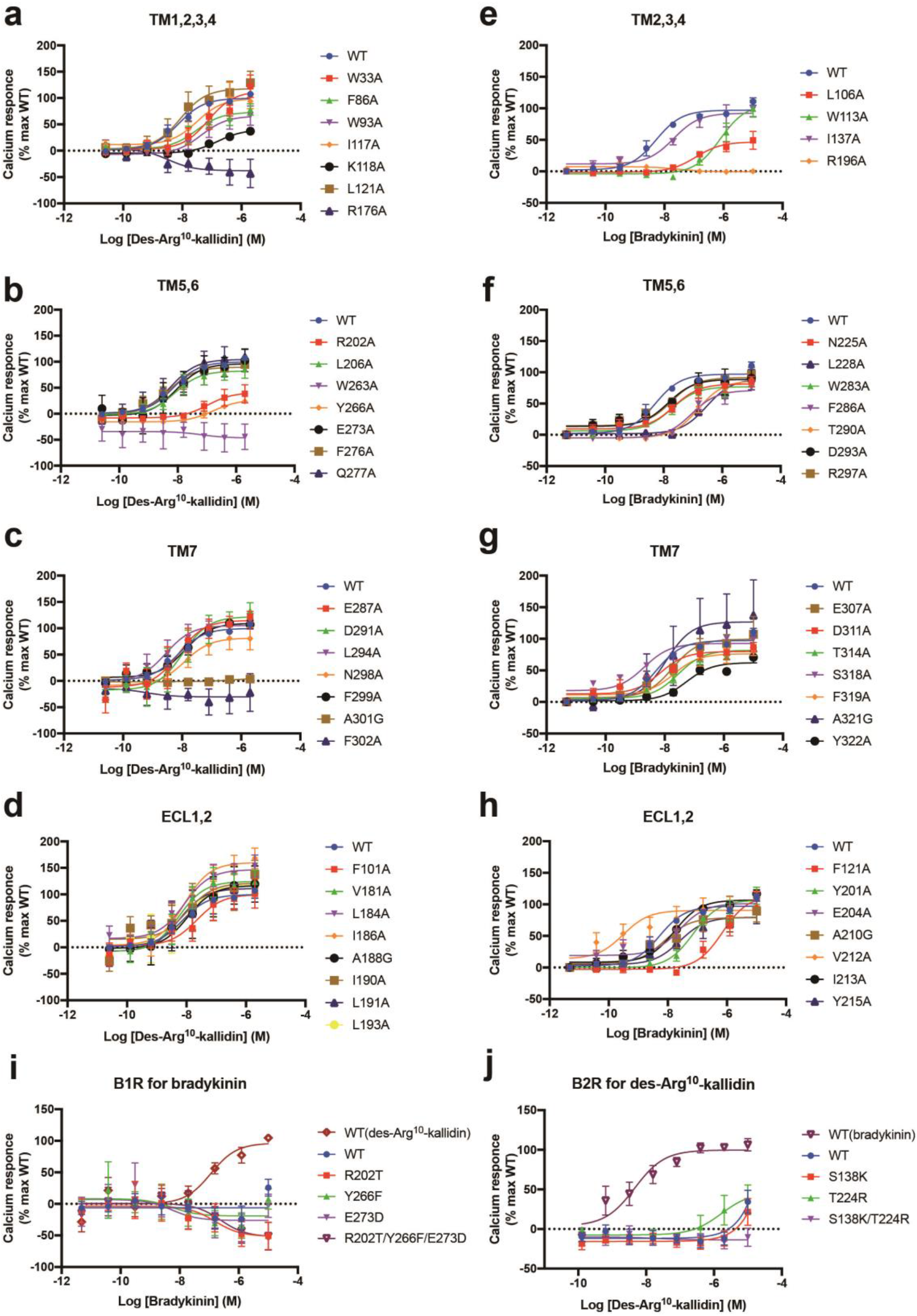
Calcium response curves of B1R and B2R. Effects of B1R mutations. (**a-d**) or B2R mutations (**f-h**) on des-Arg^10^-kallidin-or bradykinin-induced calcium mobiliazation. **i, j**, Effects of des-Arg^10^-kallidin and bradykinin on bradykinin receptors with swapped mutations. Each data point presents mean ± S.E.M. of three independent experiments. WT, wild-type.

**Extended Data Figure. 8│.**
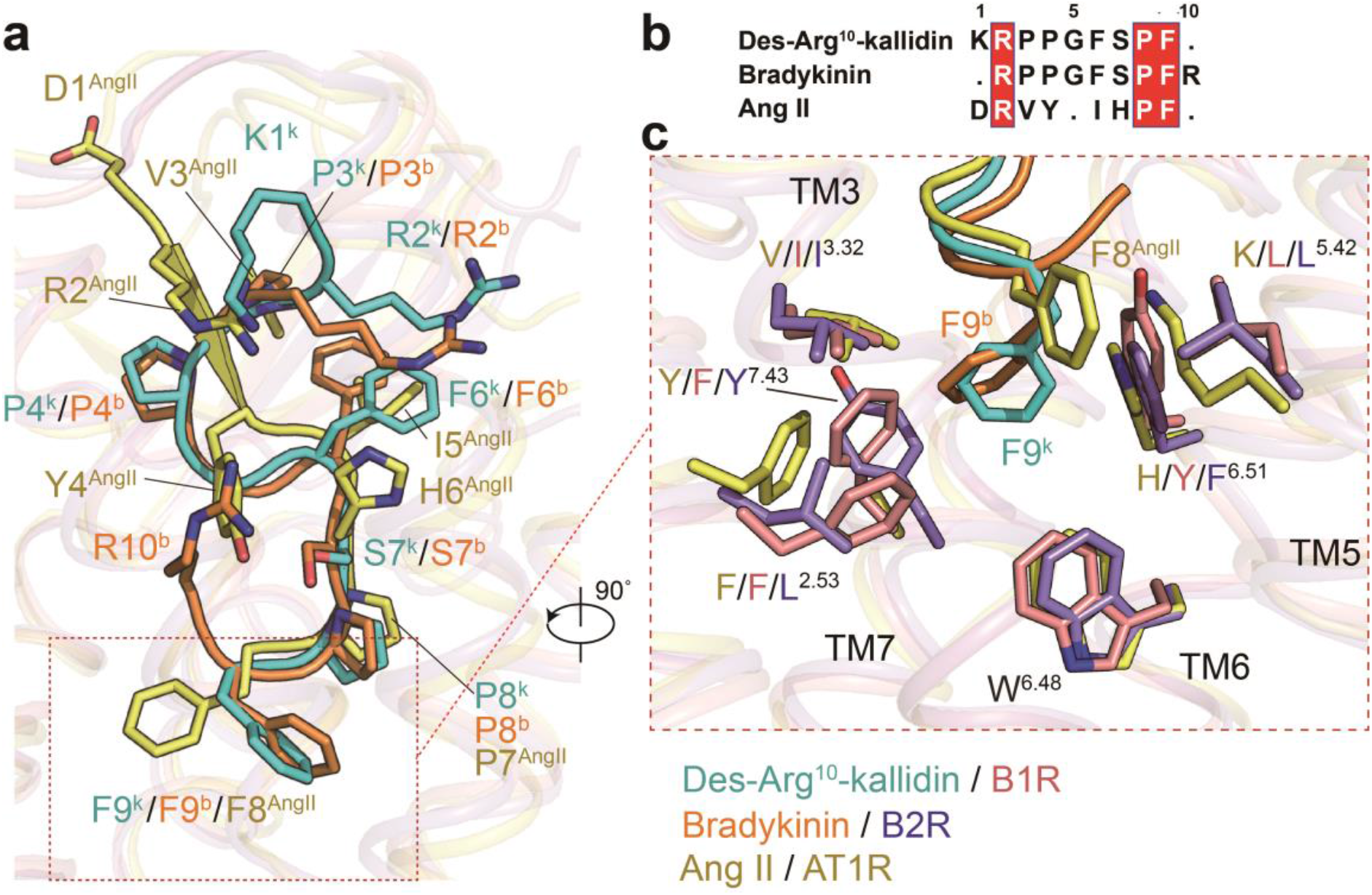
Conformational comparison of des-Arg^10^-kallidin and bradykinin with angiotensin II. **a**,**b**, The conformational comparison (**a**) and sequence alignment (**b**) of des-Arg^10^-kallidin, bradykinin, and angiotensin II (AngII). **c**, A structural comparison of conserved phenylalanine in peptides and their surrounding residues in corresponding receptors. Peptides and receptors are colored as indicated. Ang II-AT1R complex (PDB: 6OS0).

**Extended Data Figure. 9│.**
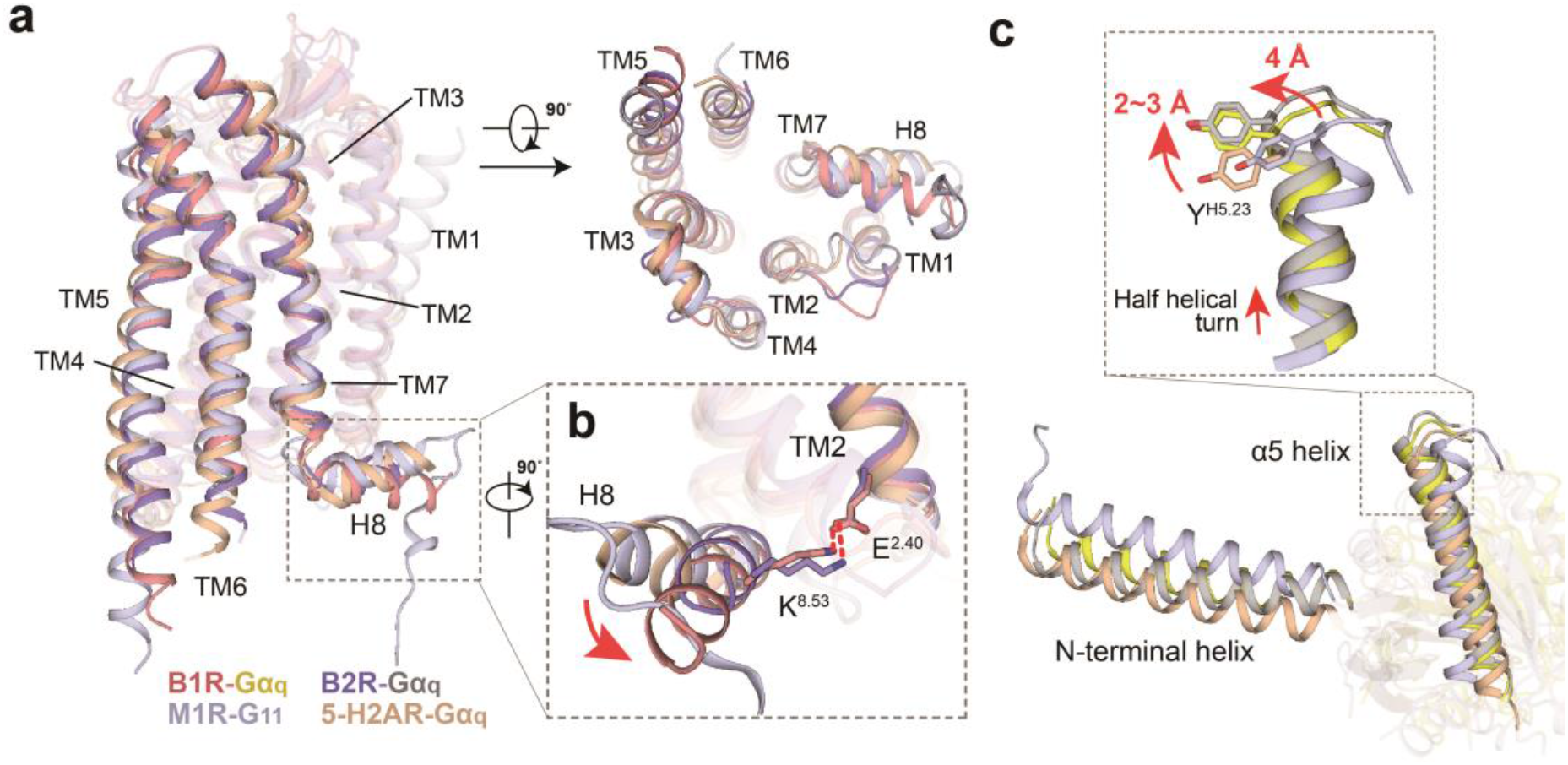
G_q_ protein-coupling of bradykinin receptors. **a**, An overall conformational comparison of two G_q_-coupled bradykinin receptors with G_q_-coupled 5-HT_2A_ R (PDB: 6WHA) and G_11_-coupled M1R (PDB: 6OIJ). **b**, A conformational comparison of the helix 8 of four G_q/11_-coupled receptors. Movement directions of helix 8 in two bradykinin receptors relative to that of 5-HT_2A_ R and M1R are indicated as red arrows. H-bonds are shown as red dashed lines. **c**, A structural comparison of α5 helix and αN of Gα_q/11_ among four G_q/11_-coupled receptor complexes. Red arrows indicate the movements of α5 helix of Gα_q_ from the bradykinin receptor-G_q_ complexes compared to 5-HT_2A_ R–G_q_ or M1R–G_11_ complexes. The complex structures were aligned by the receptors.

**Extended Data Figure. 10│.**
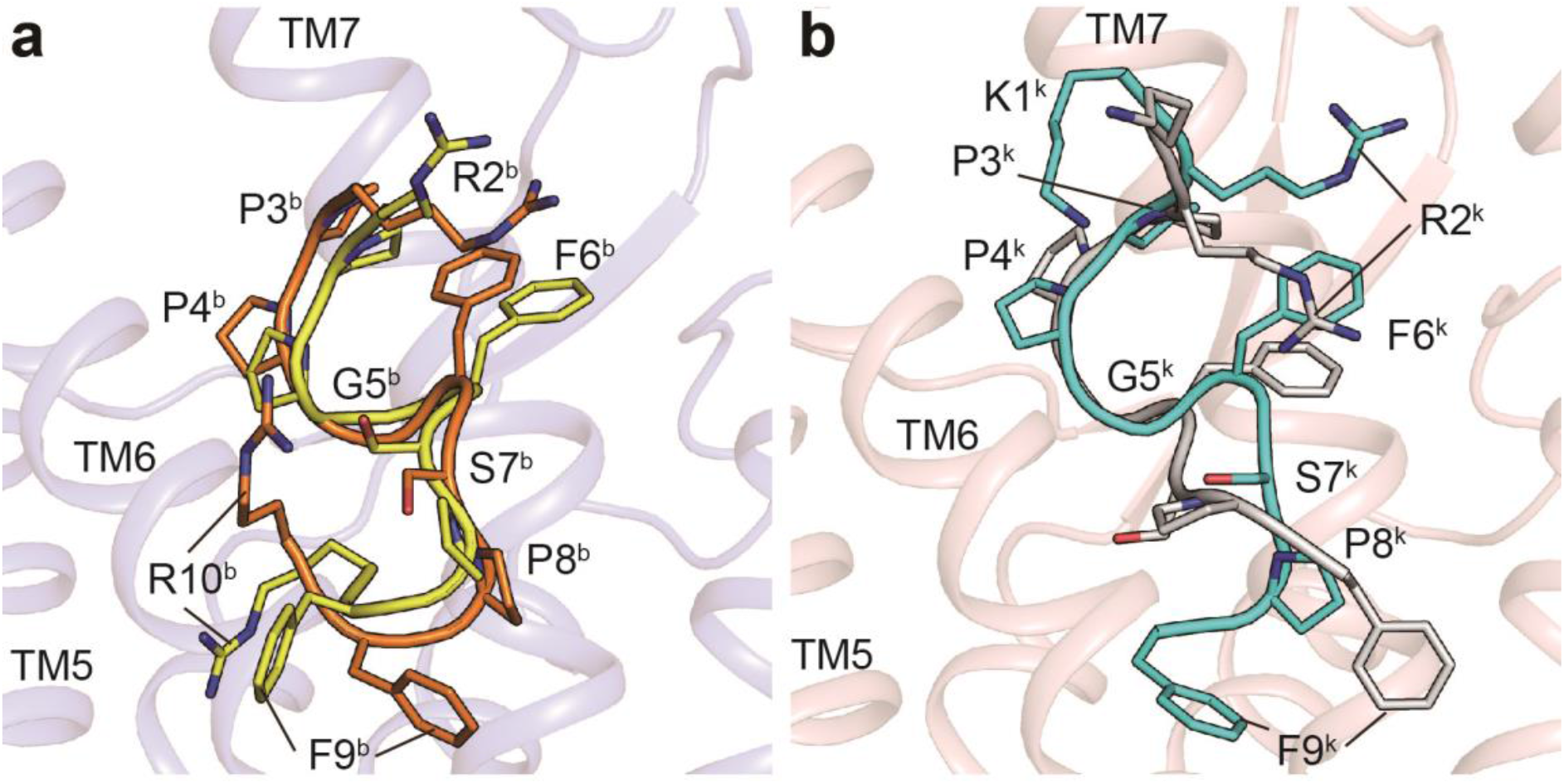
Conformational comparison of kinins in structure and in NMR model. **a**, Conformational superposition of bradykinin in B2R structure (orange) and in NMR model (yellow, PDB: 6F3V). **b**, Conformational superposition of des-Arg^10^-kallidin in B1R structure (cyan) and in NMR model (silver, PDB: 6F3Y). Kinins are shown as cartoons, and side chains are displayed as sticks.

**Extended Data Table 1 │.** Cryo-EM data collection, model refinement and validation statistics.

|  | B1R-Gq | B2R-Gq |
| --- | --- | --- |
| **Data collection and processing** |  |  |
| Magnification | 64,000 | 81,000 |
| Voltage (kV) | 300 | 300 |
| Electron exposure (e<sup>-</sup>/Å<sup>2</sup>) | 61.8 | 80 |
| Defocus range (μm) | -1.2 ~ -1.8 | -1.2 ~ -2.2 |
| Pixel size (Å) | 1.08 | 1.045 |
| Symmetry imposed | C1 | C1 |
| Initial particle projections (no.) | 3,681,755 | 3,460,328 |
| Final particle projections (no.) | 633,636 | 664,416 |
| Map resolution (Å) | 3.0 | 2.9 |
| FSC threshold | 0.143 | 0.143 |
| Map resolution range (Å) | 2.0-4.0 | 2.0-4.0 |
| **Refinement** |  |  |
| Initial model used | 6OS0 | 6JOD |
| Model resolution (Å) | 3.1 | 3.3 |
| FSC threshold | 0.5 | 0.5 |
| Map sharpening *B* factor (Å<sup>2</sup>) | -116.69 | -103.22 |
| Model composition |  |  |
| Non-hydrogen atoms | 7206 | 9004 |
| Protein residues | 928 | 1150 |
| Lipids | 0 | 0 |
| *B* factors (Å<sup>2</sup>) |  |  |
| Protein | 59.87 | 65.10 |
| Lipids | - | - |
| R.m.s. deviations |  |  |
| Bond lengths (Å) | 0.006 | 0.003 |
| Bond angles (°) | 0.904 | 0.564 |
| Validation |  |  |
| MolProbity score | 1.14 | 1.49 |
| Clashscore | 3.45 | 5.48 |
| Rotamer outliers (%) | 0.27 | 0.41 |
| Ramachandran plot |  |  |
| Favored (%) | 98.03 | 96.81 |
| Allowed (%) | 1.97 | 3.19 |
| Disallowed (%) | 0.00 | 0.00 |

**Extended Data Table 2 │.** Effects of alanine scanning mutagenesis of the residues in the binding pocket of B1R/B2R on des-Arg^10^-kallidin/bradykinin-induced calcium mobilization.

| **Residue number** | **B1R mutant** | **Calcium mobilization**<br>**pEC<sub>50</sub> ± S.E.M.** | **B2R mutant** | **Calcium mobilization**<br>**pEC<sub>50</sub> ± S.E.M.** |
| --- | --- | --- | --- | --- |
| - | WT | 8.17±0.22 | WT | 8.35±0.17 |
| 1.28 | W33A | 7.18±0.46 | W55A | N.T. |
| 2.53 | F86A | 7.52±0.27 | L106A | 6.97±0.29* |
| 2.60 | W93A | 7.29±0.22* | W113A | 6.08±0.24** |
| ECL1 | F101A | 7.60±0.26 | F121A | 6.15±0.36** |
| 3.32 | I117A | 7.45±0.46 | I137A | 7.95±0.31 |
| 3.33 | K118A | 6.78±0.32* | S138A | N.T. |
| 3.36 | L121A | 8.03±0.27 | L141A | N.T. |
| 4.64 | R176A | N.A. | R196A | N.A. |
| ECL2 | V181A | 8.27±0.14 | Y201A | 7.20±0.12** |
| ECL2 | L184A | 8.26±0.26 | E204A | 7.64±0.27 |
| ECL2 | I186A | 8.11±0.22 | V208A | N.T. |
| ECL2 | A188G | 8.02±0.43 | A210G | 7.93±0.24 |
| ECL2 | I190A | 7.56±0.79 | V212A | 9.42±0.95 |
| ECL2 | L191A | 8.39±0.45 | I213A | 7.78±0.23 |
| ECL2 | L193A | 8.35±0.66 | Y215A | 7.85±0.47 |
| 5.38 | R202A | 6.64±0.41* | T224A | N.T. |
| 5.39 | I203A | N.T. | N225A | 7.68±0.12* |
| 5.42 | L206A | 7.99±0.29 | L228A | 6.57±0.14** |
| 6.48 | W263A | N.A. | W283A | 7.75±0.12* |
| 6.51 | Y266A | 6.84±0.31* | F286A | 7.03±0.08** |
| 6.55 | A270G | N.T. | T290A | 6.80±0.14** |
| 6.58 | E273A | 8.25±0.49 | D293A | 7.90±0.25 |
| 6.61 | F276A | 8.42±0.31 | H296A | N.T. |
| 6.62 | Q277A | 8.39±0.52 | R297A | 7.84±0.38 |
| 7.28 | E287A | 8.24±0.33 | E307A | 7.99±0.40 |
| 7.32 | D291A | 8.12±0.43 | D311A | 8.22±0.24 |
| 7.35 | L294A | 8.72±0.59 | T314A | 7.54±0.13* |
| 7.39 | N298A | 7.92±0.15 | S318A | 8.52±0.28 |
| 7.40 | F299A | 8.03±0.30 | F319A | 7.87±0.28 |
| 7.42 | A301G | N.A. | A321G | 7.89±0.50 |
| 7.43 | F302A | N.A. | Y322A | 7.40±0.20* |

Data are presented as means ± S.E.M. of three independent experiments. All data were analyzed by Student *t*-test. \**P*<0.05, \*\**P*<0.01 vs. wild-type (WT). N.A., not active; N.T., not tested.

**Extended Data Table 3 │.** Effects of mutagenesis of the residues interacting with F9^k^/F9^b^ of B1R/B2R on des-Arg^10^-kallidin/bradykinin-induced calcium mobilization.

| **B1R mutant** | **Calcium mobilization**<br>**pEC<sub>50</sub> ± S.E.M.** | **B2R mutant** | **Calcium mobilization**<br>**pEC<sub>50</sub> ± S.E.M.** |
| --- | --- | --- | --- |
| WT | 7.08±0.26 | WT | 8.10±0.12 |
| K118S | N.A. | T224R | 8.03±0.42 |
| R202T | N.A. | F286Y | 7.17±0.19* |
| K118S/ R202T | N.A. | D293E | 7.50±0.23 |
| - | - | T224R/F286Y/D293E | 5.45±0.21*** |

Data are presented as means ± S.E.M. of three independent experiments. All data were analyzed by Student's *t*-test. \**P*<0.05, \*\*\**P*<0.001 vs. wild-type (WT). N.A., not active.

